# The time dimension matters: Improving mode of action classification with live-cell imaging

**DOI:** 10.1101/2025.04.22.649936

**Authors:** Edvin Forsgren, Jonne Rietdijk, David Holmberg, Julia Juneblad, Bianca Migliori, Martin M. Johansson, Jordi Carreras-Puigvert, Johan Trygg, Gillian Lovell, Ola Spjuth, Pär Jonsson

## Abstract

Morphological profiling is a common approach to investigate the modes of action (MOAs) of compounds. Most methods rely on fixed-cell assays, which provide only a single snapshot at a predefined time point and overlook the dynamic nature of cellular responses. In contrast, live-cell imaging tracks responses over time, offering deeper insight into compound-specific effects and mechanisms; however, time-series analysis of image data remains challenging due to limited analytical tools.

We present Live Cell Temporal Profiling (LCTP), a workflow for morphological profiling of label-free live-cell time series data that yields interpretable, biologically relevant results. We showcase LCTP in an MOA classification study using label-free data. The workflow integrates established deep-learning components, cell segmentation, live/dead classification, and single-cell feature extraction, with data-driven models to capture MOA-specific temporal phenotypes and produce time-resolved profiles that can be compared across compounds and cell lines.

We assess MOA classification performance using double-blinded cross-validation simulating a real-world screening scenario. LCTP significantly improves MOA classification over single–time point analysis, consistently across both cell lines used in the study. Time-resolved phenotypic modeling reveals transient, sustained, and delayed responses, clarifying compound-specific temporal effects and mechanisms across MOAs.

The presented workflow is modular: each step removes irrelevant information, enriching signal, and enabling straightforward updates as technologies evolve and as new technologies become available, while supporting reuse across studies broadly. We believe LCTP adds substantial value to high-throughput compound screening, showing that live-cell imaging combined with this workflow yields informative visualizations of temporal effects and improved MOA classification.

## 1. Introduction

Time-series imaging captures the temporal, complex dynamics of biological systems. Yet, extracting meaningful insights from such high-dimensional, cellular data remains a significant challenge in modern drug discovery[1, 2]. The ability to analyse how cells respond to compounds over time, rather than at arbitrary endpoints, represents a critical step for understanding therapeutic modes of action (MOA) and predicting clinical outcomes [3].

One powerful tool for studying the MOA of drugs involves capturing and analysing cellular phenotypic changes in response to treatment - an approach known as image-based morphological profiling [4, 5]. By leveraging high-content imaging techniques such as the Cell Painting assay [6, 7, 8], researchers can generate rich, unbiased datasets that reveal how different compounds alter cellular structures and organelle organization [7]. These morphological fingerprints can be compared across a reference database of known drug effects, enabling the identification of shared phenotypic signatures to infer MOA and support virtual screening [9].

Today, morphological profiling is typically performed as a single end point assay, where compounds are applied for a set period, such as 48 hours, followed by cell fixation and staining using fluorescent dyes or antibodies. This requires complex staining protocols, potentially introducing technical variations due to factors such as reagent lots, processing times, equipment calibration, and experimental platforms, leading to batch effects between plates and laboratories. Additionally, an endpoint assay only provides a static snapshot of cellular states, potentially missing important effects that occurred earlier or have not yet happened and thus missing important time-dependent dynamics.

Biological processes are inherently dynamic. Cells continuously respond to their environment, activate complex signalling cascades, undergo division, differentiate, and ultimately die. These processes occur across varying timescales, from seconds for ion channel activity to hours for gene expression changes to days for phenotypic transformations [10]. Capturing this temporal complexity is fundamental to understanding both normal physiology and disease states. Living cells represent dynamic, adaptive systems rather than static entities, making time-resolved analysis essential for accurate biological insights.

In cancer drug development, temporal monitoring of drug-treated cells provides insights into whether cells undergo senescence, develop alternative survival pathways, or initiate a programmed cell death such as apoptosis which are key factors for designing long-lasting cancer treatments [3]. Live-cell imaging has revealed new insights into tumour and T cell behaviour, leading to the discovery of novel druggable targets to optimize engineered T cell function for improved tumour targeting [11], and identified effective drug combinations for treating heterogeneous tumour populations [12, 13].

To capture temporal dynamics in live cells, researchers commonly use permeable, live-cell-compatible stains. However, these stains can become unstable over time, and repeated imaging exposures may lead to phototoxicity due to the damaging fluorescent light. To minimize these effects, assay conditions must be carefully optimized, leading to a trade-off between assay duration, signal quality, and cell health [14].

To circumvent challenges associated with stain instability and phototoxicity, whilst also reducing cost and labour, label-free imaging offers a compelling alternative [15]. It preserves native cellular states and avoids artefacts introduced by labelling. When combined with AI-driven segmentation and feature extraction, label-free imaging provides a powerful, non-invasive approach for live-cell analysis. While label-free live-cell datasets exist [16], focus has been mostly on segmentation or cell tracking [17]. Many studies rely on a limited set of handcrafted features for characterizing cellular phenotypes, or on quantitative phase imaging (QPI), which lacks the fine subcellular detail captured by phase contrast microscopy [18, 19]. Analysis on cellular phenotypes with high-dimensional features from label-free live-cell time series data has not been widely explored.

AI and deep learning are revolutionizing the possibilities of morphological profiling by enabling the extraction of complex, high-dimensional features from imaging data with unprecedented accuracy and scalability [4]. With these new technologies, high-dimensional features can be readily extracted from label-free cell imaging. And unlike traditional analysis methods that rely on predefined features, deep learning models can automatically learn intricate phenotypic patterns, improving sensitivity in detecting subtle cellular changes[20, 21].

Yet, in practice, analysis of live-cell morphological profiling remains largely time-point centric. Common workflows, whether based on dimensionality reduction, clustering, or supervised classifiers, are typically applied independently at each time point, yielding a series of disconnected snapshots and a lot of manual interpretation. A time-series perspective instead treats phenotypes as trajectories in high-dimensional space which we can leverage to understand biology better in terms of both timing and magnitude of change. In other data-rich monitoring domains, such as bioprocessing, multivariate time-resolved modelling and trajectory-based signatures are standard for extracting robust signal from noisy, high-dimensional measurements; bringing the same principles to live-cell phenotyping should similarly improve sensitivity to weak or transient effects. For label-free imaging, where phenotypic effects may be subtle, the ability to form coherent temporal profiles is particularly valuable. Shifting from isolated end points to time series data can provide a more faithful account of cellular responses and give researchers a more nuanced understanding of complex biological events.

In this paper, we demonstrate the value of time series analysis compared to single time point analysis by comparing the accuracy of MOA classification on two separate cell lines, A549-WT and U2OS-WT. To do this, we developed a multi-model workflow to handle temporal data where we utilize deep learning to convert time series of label-free live-cell images into interpretable profiles showcasing compounds dynamic effects over time, Fig 1. We refer to this as Live Cell Temporal Profiling (LCTP). LCTP consists of pre-trained deep learning models for cell segmentation, live/dead classification, and single cell feature extraction followed by supervised time-dependent phenotypic modelling. Our results reveal that LCTP provides valuable insights into the temporal dynamics of compound effects and significantly enhances accuracy in MOA classification compared to a single time point analysis.

**Figure 1.**
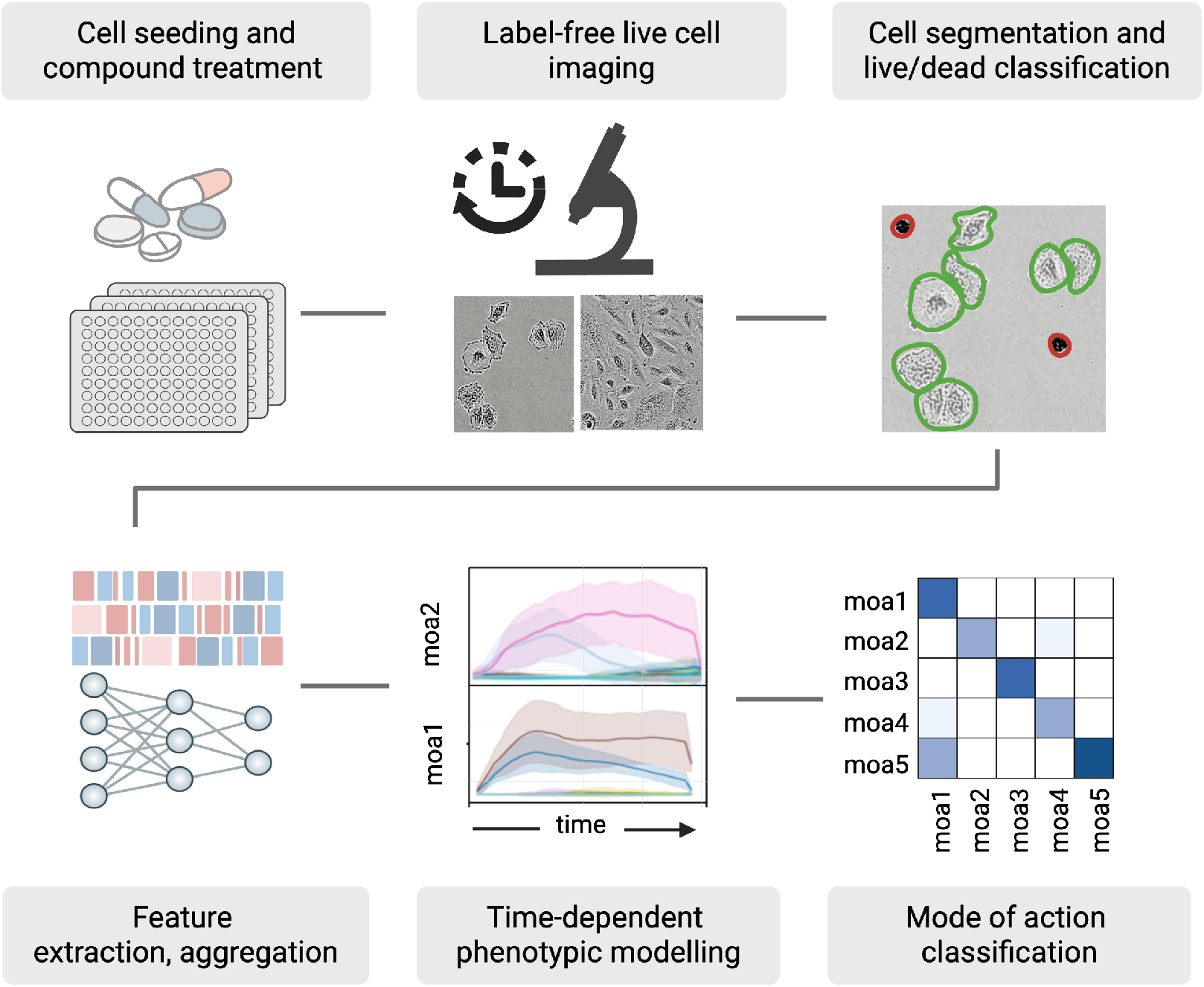
Schematic workflow of the time-dependent phenotypic profiling approach. Cells are seeded and treated with compounds in a multi-well plate format, followed by live-cell imaging at multiple time points. Images are then used to segment single cells and to classify live and dead cells. Single-cell features are extracted using a feature extraction model, after which the features are aggregated to median well features. These features are used in Live Cell Temporal Profiling (LCTP), a supervised, time-dependent phenotypic modelling approach to capture dynamic cellular responses. The resulting predictions form time-dependent signatures that provide insights into temporal effects and are used for MOA classification.

## 2. Materials and Methods

### 2.1. Cell assay and live-cell imaging

Prior to assay, compound plates were thawed for 30 minutes at room temperature in a sterile biological safety cabinet. Cells (A549-WT, U2-OS-WT; Supplementary Table 1) were cultured in T75 cell culture flasks and harvested on the day of assay by aspiration of media, washing with 10 mL PBS, and addition of 1.5 mL Trypsin-EDTA solution. Once detached, cells were suspended in fresh media and counted (Invitrogen Countess 3 FL). Cell suspensions were formed in 50 mL centrifugation tubes (2.5 E+04 cells/mL, 13 mL per plate). After mixing well, the cell suspension was added into the 384-well compound plate in 30 µL/well volume using a manual multichannel pipette. Plates were centrifuged for 1 minute at 100xg to align cells at the bottom of the well, then placed into an Incucyte^®^ SX5 Live Cell Analysis System equipped with a Green/Orange/NIR optical module. Images were acquired using Adherent Cell-by-Cell acquisition at 20x magnification every 2h for 3 days, with a single image site per well. Images were analysed using AI Cell Health Analysis^®^ (Supplementary Table 2). which provides individual cell segmentations and classification as live or dead[22, 23]

### 2.2. Compound treatments

We selected compounds belonging to six diverse MOA classes that we intended to range from subtle to clear changes in morphology and are expected to be either fast-acting or slow-acting. Specifically, we included AKT inhibitor (AKT), CDK inhibitor (CDK), HDAC inhibitor (HDAC), MAP kinase inhibitor (MAPK), PARP inhibitor (PARP) and tubulin polymerization inhibitor (TUB). Of these, we expected TUB to be easy to separate from others and untreated cells due to their distinct morphology (Fig 2E) and PARP to be the most challenging due to subtle morphological changes that accumulate over time due to DNA damage and thus hard to separate from untreated cells [24]. For each of these MOAs we selected twelve compounds from the SPECS repurposing library. To account for differences in compound potency and toxicity, we used internal data from compounds tested at 10 µM to ensure detectable effects without inducing toxicity (data not published).

**Figure 2.**
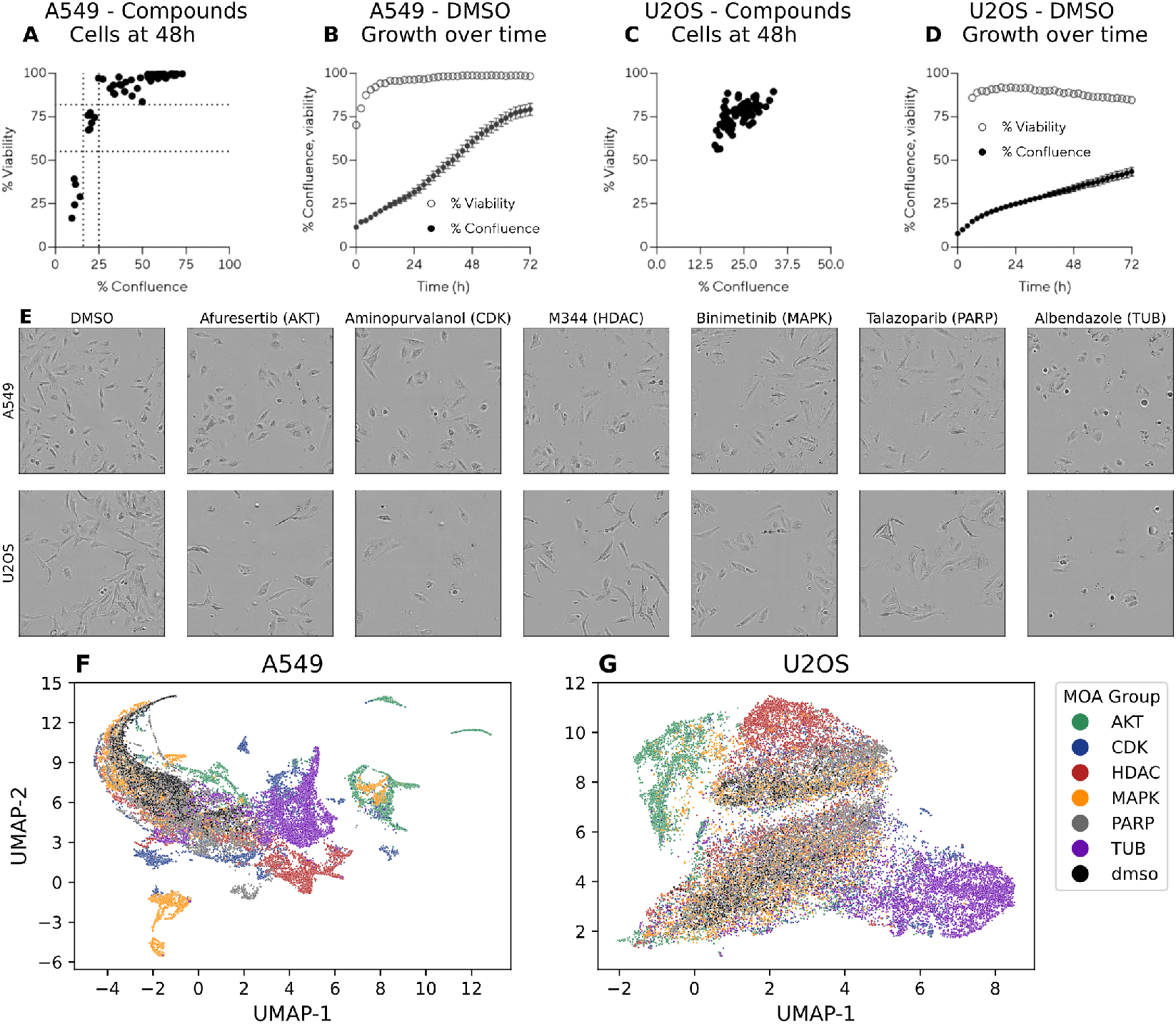
Scatter plots, A549 in A and U2OS in C, indicates the mean % confluence and % viability values of each compound at 48h. Time series plot shows the % confluence and % viability of A549 cells in B and U2OS cells in D in the negative control (DMSO) wells over time. The mean value comprising all negative control wells from all plates is displayed and error bars represent standard deviation. In E, phase contrast images show negative control and compound-treated A549 and U2OS cells exemplifying each mode of action at 48h. In F and G, UMAPs for A549 and U2OS cells including data from every other hour from 24h to 72h.

Each compound was tested in fifteen replicates, consisting of five technical replicates (i.e., compound treatments on the same plate) and three biological replicates (i.e., treatments on different plates). Each plate included fifteen wells containing only the solvent, DMSO (Dimethyl Sulfoxide), as negative controls. Each of the six mircroplates was imaged at different occasions to simulate a real world setting.

The plate layout for the six microplates, three per cell line, was performed using PLAID (Plate Layouts using Artificial Intelligence Design [25], a constraint-programming-based method designed to minimize unwanted well-position and plate effects. Compound handling was performed by the SciLifeLab Compound Center (CBCS, Solna, Stockholm). Briefly, chemicals were solubilized in either DMSO or water at stock concentrations between 2 and 10 mM, as specified in Supplementary Table 6. Compounds were dispensed in nanoliter volumes using the Echo liquid handler into 384-well PhenoPlates (Revvity, cat. no. 6057302) and stored at −20°C prior to experimentation.

### 2.3. Characterisation of live cell assay

The live-cell imaging data was used to examine the effect of these compounds on A549 and U2OS cells using the AI Cell Health Analysis^®^ module [23, 22]. Untreated (DMSO) A549 cells displayed high viability throughout the three-day time frame of the assay, and proliferation was observed as the confluence increased from ~12% to over 75%, Fig 2B. The single time point (48h) scatter plots, Fig 2A, indicate that a small number of compounds strongly affected A549 viability, showing a cluster of compounds in the bottom left quadrant that decreased viability, inducing cell death. Another cluster of compounds can be observed in which cells have reduced confluence (<25%) but maintain viability in the range of 60-80%, suggesting a cytostatic mechanism and the possibility of cell cycle arrest. The remaining compounds retain >80% viability in most cases with a range of effects on cell growth indicated by confluence values ranging from 25 to 80%. Fig 2E displays negative control A549 cells with a normal healthy adherent cell morphology, and then the effect of treatments showing one example from each MOA. Afuresertib (AKT) subtly altered the shape, with an increase in circularity and texture. Albendazole (TUB) generated ruffling in the plasma membrane boundary, with a higher proportion of visible mitotic cells. Aminopurvalanol (CDK) treated cells displayed an increase in contrast around the nuclear envelope. Binimetinib (MAPK) increased cell area with prominent focal adhesion points. M344 (HDAC) and Talazoparib (PARP) both increased cell area; with M344-treated cells appearing ruffled at the boundary while Talazoparib-treated cells maintained a more circular appearance.

Untreated (DMSO negative control) U2OS cells also displayed high viability throughout the time course of the assay, however, the rate of proliferation was lower than that of A549, reaching just under 50% by the end of three days as displayed in Fig 2D. In contrast to A549 cells, the single time point (48h) scatter plot, Fig 2C, indicates that fewer compounds affected U2OS viability. Fig 2E shows that untreated U2OS cells display a heterogeneous, adherent morphology with lower contrast relative to A549 cells. Treatment with Afursertib (AKT) induced a very subtle change in morphology with a slight increase in cytosolic texture which can indicate stress. Albendazole (TUB) treatment increased the proportion of cells in mitosis with remaining cells appearing more rounded. Cells treated with Aminopurvalanol (CDK) appeared enlarged, flattened, and rounded while those treated with Binimetinib (MAPK) and Talazoparib (PARP) experienced increased area and roundness. HDAC inhibitor M344 (HDAC) resulted in more cells appearing elongated and with higher contrast.

In Fig 2F and G, scatter plots display unsupervised UMAP [26] embeddings (n_neighbours=25, min_dist=0.1, metric=“euclidean”) of cell morphology features extracted using DINOv2-ViT-B14 (see section *Pre-trained models: Segmentation and feature extraction*) from wells containing A549 and U2OS cells, coloured by MOA group. The data span 24h to 72h, with each point representing a well at a specific time point. For A549, Fig 2F, distinct clusters of MOA groups emerge, though many MOAs remain heterogeneous, forming scattered clusters, for example, AKT (green), CDK (blue), and MAPK (yellow). In contrast U2OS, Fig 2G, show less distinct clustering overall, with only a few recognizable groups such as TUB (purple), clusters of AKT, and HDAC (red). UMAP visualizations provide an overview of the data, and colouring points by known metadata helps reveal patterns, both desired (MOA/compound clusters) and undesired (batch effects). However, while these embeddings highlight structure, interpreting the specific effects of different MOA groups on cells remains challenging since unsupervised methods such as UMAPs are designed to capture overall variance of the dataset and not specific differences between groups.

### 2.4. Model and training details

The ANNs used in this work for LCTP are fully connected deep learning architectures implemented to predict soft MOA labels from 768 deep learning features (explained in more detail in section *Data analytical workflow*). For these deep learning features, the model takes the vector of 768 features as input followed by two hidden layers, consisting of 512 and 256 neurons, and an output layer with six neurons, one for each MOA in our dataset. The network use LeakyReLU activation functions in the hidden layers, and a high dropout rate of 0.5 to reduce overfitting. The output layer uses a linear activation (no softmax), allowing outputs to range beyond [0, 1]. This produces ‘soft’ equivalence scores (Eq. scores) [27] that capture graded MOA similarity rather than forcing discrete classification. During training, target labels are binary (0, 1), but predictions can fall outside this range, with higher values indicating stronger MOA membership. The model uses MSE Loss and the AdamW optimizer with a learning rate of 0.0004. The model was implemented in pytorch [28] and was trained for 200 epochs with no early stopping. For our time series approach the full CV scheme takes ~5 minutes to run per cell line with parallelization on one NVIDIA GeForce RTX 3090 and a batch size of 64.

## 3. Live Cell Temporal Profiling - Data Analytical Workflow

Live Cell Temporal Profiling (LCTP) is a data analytical workflow for analysing time series imaging data to extract interpretable information from this complex, temporal, unstructured information. The extracted information is then used to identify time trends of compounds treatment as well as MOA classification. In this section we go into detail of the different steps that constitutes LCTP.

### 3.1. Pre-trained models: Segmentation and feature extraction

Single-cell segmentation masks, derived from the label-free AI Cell Health Analysis^®^ [22, 23], were employed to extract individual cells. These cells were cropped and zero-padded at their native resolution to a fixed size of 224 × 224 pixels. Cells exceeding this size or located at image boundaries were excluded, leading to the removal of approximately 18% of the A549-WT cells and 21% of the U2OS-WT. Out of these, only 0.4% and 1.7% respectively were removed due to size and the rest were edge cells. A Chi-squared test (*p* = 0.997) confirmed no significant variation in exclusion proportions across the six MOA groups, supporting the null hypothesis *H*_0_ of MOA-independent exclusion. We chose to use fixed-size, no-resize crops to (i) avoid interpolation artifacts introduced by resizing, (ii) prevent artificial changes in apparent cell scale that could shift extracted features, and (iii) eliminate biases from variable input dimensions the feature extractor, which alter the patch grid and the amount of surrounding context the model sees. We used 224×224 because it matches the DINO ViT-B/16 pretraining resolution, using the model’s native crop size also avoids positional-embedding interpolation and scale mismatches, which typically yields more consistent, comparable features than markedly different input sizes. By doing so with fixed-size, no-resize crops we get more comparable features and ensure that the features reflect true biological differences.

The processed cell crops were then passed through a pre-trained DINOv2-ViT-B14 model, extracting 768 features per single cell [29, 30, 31] from which median well-aggregated features were computed for each time point. Using the cell health labels, i.e. living or dead labels, from the AI Cell Health analysis we created two feature sets; one based on both living and dead cells, and one based solely on living cells. We hypothesized that restricting the analysis to living cells would capture more relevant information about cellular condition and function, rather than reflecting post-mortem morphological changes. The remainder of this paper focuses on the living cell feature set. The results including dead cells are presented in the supplementary information for comparison, which supported our hypothesis. Each plate was normalized separately using a z-score transformation:

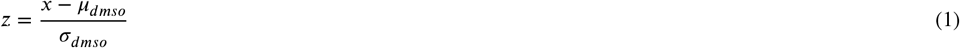

where *μ*_*dmso*_ and *σ*_*dmso*_ are the mean and standard deviation computed from negative control (DMSO) wells on the same plate. This per-plate normalization accounts for batch effects while preserving the biological signal. These processed outputs constitute the DINO feature set.

### 3.2. Data-driven models: Artificial Neural Networks (ANNs)

Based on these median well features, we trained two ANNs: one for capturing early MOA-specific phenotypes (6-14 hours) and another for late phenotypes (46-54 hours). Each ANN was trained using data within its designated time window, along with negative control data from both windows. These models utilize median well features as inputs, with each sample representing a specific well and time point. The ANNs have six outputs, each corresponding to a distinct MOA group. During training, wells exposed to compounds associated with a given MOA were assigned an output value of one for the corresponding MOA, while the remaining outputs were set to zero.

The models employ a soft classifier approach, which assigns values on a continuous scale rather than discrete class labels. This approach introduces flexibility, allowing predictions to fall below zero or above one, ensuring that compounds with weaker but similar effects are not strictly classified as belonging to a group but rather assigned intermediate values.

### 3.3. Eq. scores and Eq. profiles

Once trained, the ANN models were applied to unseen samples, generating predictions for each MOA group across all wells and time points. These predictions are referred to as Equivalence Scores (Eq. scores) [27], with one Eq. score assigned per MOA group. These are designed to convert abstract, high-dimensional features to interpretable scores. Eq. scores from all time points for a given well form Eq. profiles, which serve as temporal signatures for MOA classification (see Fig 3).

**Figure 3.**
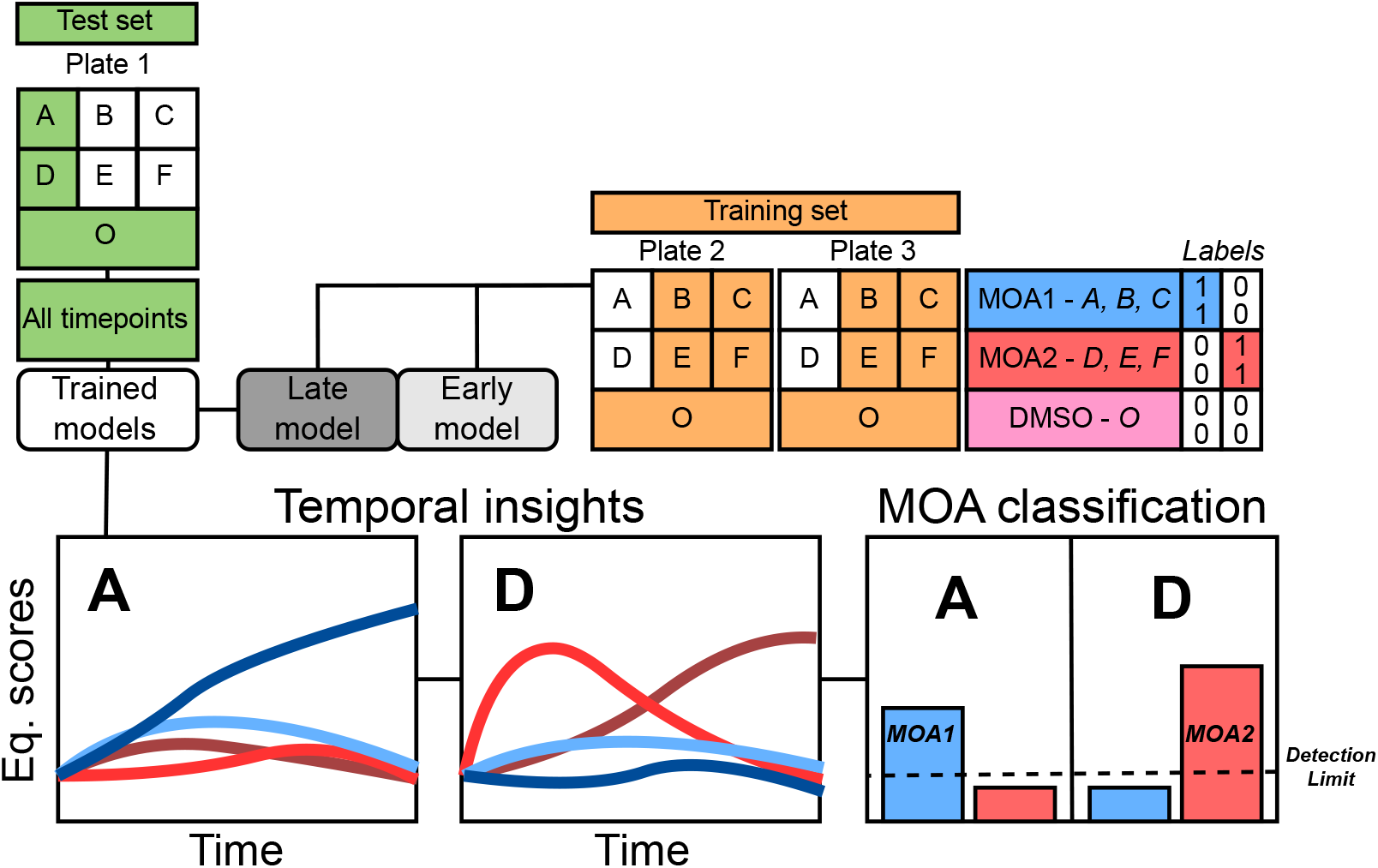
Illustration of one round in a 9-round double blinded (plate-compound) cross-validation (CV) used in the data analysis workflow. This schematic uses a smaller study design solely to illustrate the CV setup. Compounds A, B, and C represent MOA group 1; D, E, and F represent MOA group 2; O denotes negative DMSO controls. Green indicates the held-out test set for the current CV round; orange indicates the training set; white indicates data excluded from the current CV round. Blue, red, and pink label MOA1, MOA2, and DMSO, respectively. In the round illustrated, all held-out samples are from plate 1: A, D, and O (DMSO) are predicted as the test set. The next two rounds on plate 1 hold out B, E, and O, and then C, F, and O, respectively. The same three-round scheme is repeated for plates 2 and 3, yielding nine CV rounds in total. In each CV round, two models are fitted, an early model (hours 6–14) and a late model (hours 46–54), using only data from plates and compounds outside the held-out predictions (e.g., in the first round, B, C, E, F, and O from plates 2 and 3 are used for training). The held-out plate is then predicted across all time points (hours 0–72), producing four Eq. scores per time point (early and late for each MOA). For each MOA, the early and late Eq. scores are smoothed over time to mitigate biological variability and then summed; the MOA with the maximum sum is assigned. Eq. scores from negative controls (O) establish a per-MOA detection limit to filter false positives and increase confidence in the predictions.

### 3.4. Smoothing with exponentially weighted moving average

To leverage temporal information while mitigating random fluctuations, Eq. profiles were smoothed using Exponentially Weighted Moving Average (EWMA) [32]. This smoothing technique enhances the robustness of MOA classification by reducing noise and emphasizing true temporal trends.

### 3.5. Method evaluation: Cross-validation procedure

For method evaluation, a double-blind cross-validation (CV) procedure was implemented to simulate real-world conditions and prevent data leakage, see Fig 3. The CV process was structured as follows:

- In each CV round, training and test sets were created. The training set comprised data from two plates, excluding wells treated with a set of six compounds (one per MOA group).
- The test set included all wells treated with these six compounds on the third plate, which was omitted from training.
- Each plate underwent 12 CV rounds, corresponding to the 12 available compounds per MOA.
- Given three plates per cell line, this resulted in a total of 36 CV rounds per cell line.

For the time series approach, where separate early and late models were trained, this led to a total of 72 models being trained per cell line. During each CV round, predictions for six MOA-specific Eq. scores were generated across all time points (0-72 hours) in the unseen test set. This resulted in two Eq. scores per MOA per well and time point - one from the early model and one from the late model. These Eq. scores were then smoothed using EWMA, producing refined Eq. profiles that capture MOA-related temporal dynamics.

For the single time point analysis, the same CV scheme was employed, but only a single model per round was trained, resulting in 36 models per cell line. Predictions were generated solely at the 48-hour time point, a commonly used endpoint in morphological profiling assays [6].

### 3.6. MOA classification: Avoiding false positives

To prevent the misclassification of compound-treated wells that do not significantly deviate from negative controls, detection limits were established. These thresholds ensure that compounds at suboptimal concentrations, which do not induce detectable morphological changes, are not falsely classified as exhibiting MOA-related effects.

Detection limits were determined from Eq. scores of negative control wells in the CV scheme (Fig 3). They were computed based on the 95th quantile of the sum of the integrals of Eq. profiles (both early and late) for each MOA group, referred to as Eq. sum. A similar approach was applied to the single time point analysis, where detection limits were based on the average plus two standard deviations of MOA-specific Eq. scores for negative controls.

Once detection limits were established, the Eq. sum for each compound-treated well was computed by integrating and summing the early and late Eq. profiles. If the Eq. sum exceeded the detection threshold, the MOA group with the highest Eq. sum was assigned as the predicted label. This filtering mechanism ensured that only significantly altered wells were classified into one of the six MOA groups, reducing false positive rates. If the Eq. sum did not pass the detection limit, it was classified as a negative control, i.e. DMSO. Notably, detection limits varied across MOA groups, reflecting differences in morphological response intensities.

## 4. Results and Discussion

Here, we present the results from LCTP where we leverage time series live-cell imaging to generate interpretable profiles that capture the dynamic effects of compounds over time. We found that examining the full time series data of cell treatments with LCTP results in a significantly higher accuracy for MOA classification compared to analysing a single time point.

### 4.1. Temporal insights

Because each compound has a unique dose-response relationship, its effective concentration range varies. When screening thousands of compounds at a single concentration, there is no guarantee that every compound will be tested at a level where its effects are detectable at the same single time point. However, by capturing the full time series, early rapid responses to a specific MOA can still be identified, even if the cells ultimately undergo death. This allows for correlation between fast-acting, high-efficacy compounds and slower responses from compounds with the same MOA but lower potency by comparing different time points.

In Fig 4, the Eq. Profiles for three example compounds from the AKT, TUB, and HDAC MOA groups are shown. Each plot contains 12 lines, representing early and late prediction for each MOA group, as indicated in the legend. However, most of them are not visible since the predictions are close to 0. Each line represents the mean CV predicted MOA Eq. score over time, i.e. the mean MOA Eq. profile, of 15 replicates across 3 plates, with a confidence interval of one standard deviation. As a result of our CV scheme, the predictions of these 15 replicates are generated by six models, one trained on early time points (6-14 hours) and one trained on late time points (46-54 hours) per plate. Lighter colours denote early model predictions, while darker colours indicate late model predictions. These two time windows are also visualized. Worth noting is that the time windows are only a small part of the full dataset but both early and late models predict observations from all time points. Importantly, all predictions shown here and forth are “pure” CV predictions, i.e. predictions of observations not seen by the model during training.

**Figure 4.**
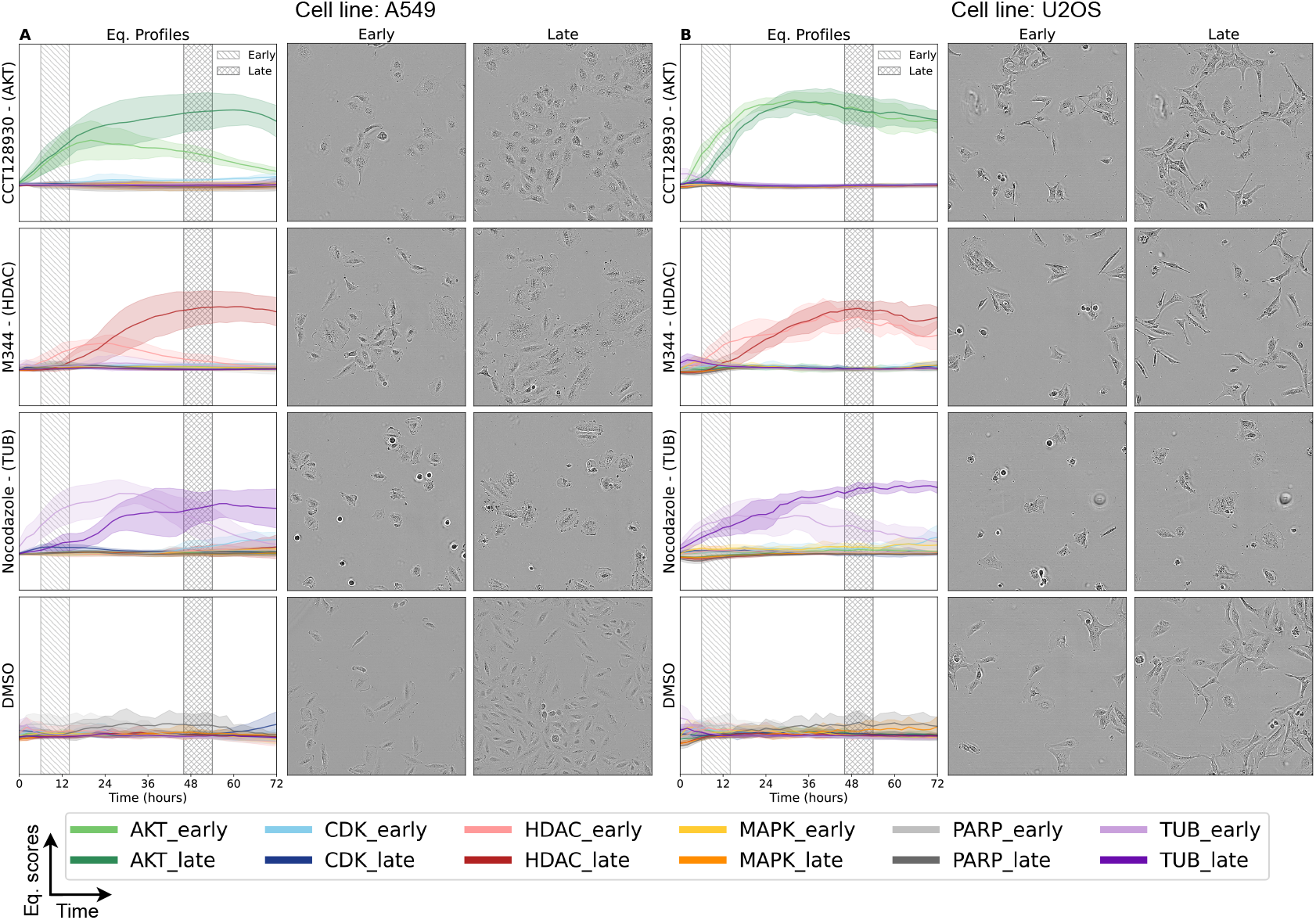
Eq. profiles and live-cell phase contrast images of A549 cells (A) and U2OS cells (B) at 8h and 48h. The first row presents data for CCT128930 (an AKT inhibitor), the second for M344 (an HDAC inhibitor), the third for Nocodazole (a TUB inhibitor), and the fourth for negative controls, DMSO, i.e. untreated cells.

Predicting across all time points can also reveal that compounds with certain MOAs may induce a transient or non-linear effect on the cells while others have a rapid and linear response. Such a response is seen for the AKT group which displays a rapid response which is sustained throughout the remainder of the time series, resulting in similar Eq. profiles for both early and late model predictions (Fig 4). Noticeably for CCT128930 (AKT), the live-cell images from 8h and 48h share similar morphologies although the confluence has increased at the later time point, confirming what can be seen in the Eq. profiles for both cell lines. This is consistent with previously observed responses to AKT inhibitors, where the AKT signalling pathway is activated within minutes of inhibitor exposure [33].

A different example are the HDAC inhibitors that for A549 cells exhibit a low early Eq. profile but with a clear late profile. Because HDAC inhibitors are believed to regulate transcription through epigenetic modifications, which typically take time to manifest [34], it is expected to see such delayed effects. For U2OS cells, the profiles show both clear early and late Eq. profiles showcasing a clear difference between the two cell lines.

Similarly to HDAC inhibitors, PARP inhibitors, which prevent the repair of single-strand DNA breaks [24], have only a very late onset and show the lowest classification accuracy (Figs 5, 6, and 7). We hypothesize that their effects become apparent only after mutations accumulate over time, possibly explaining the delayed effects observed and might even be more evident at later time points than acquired in this dataset.

**Figure 5.**
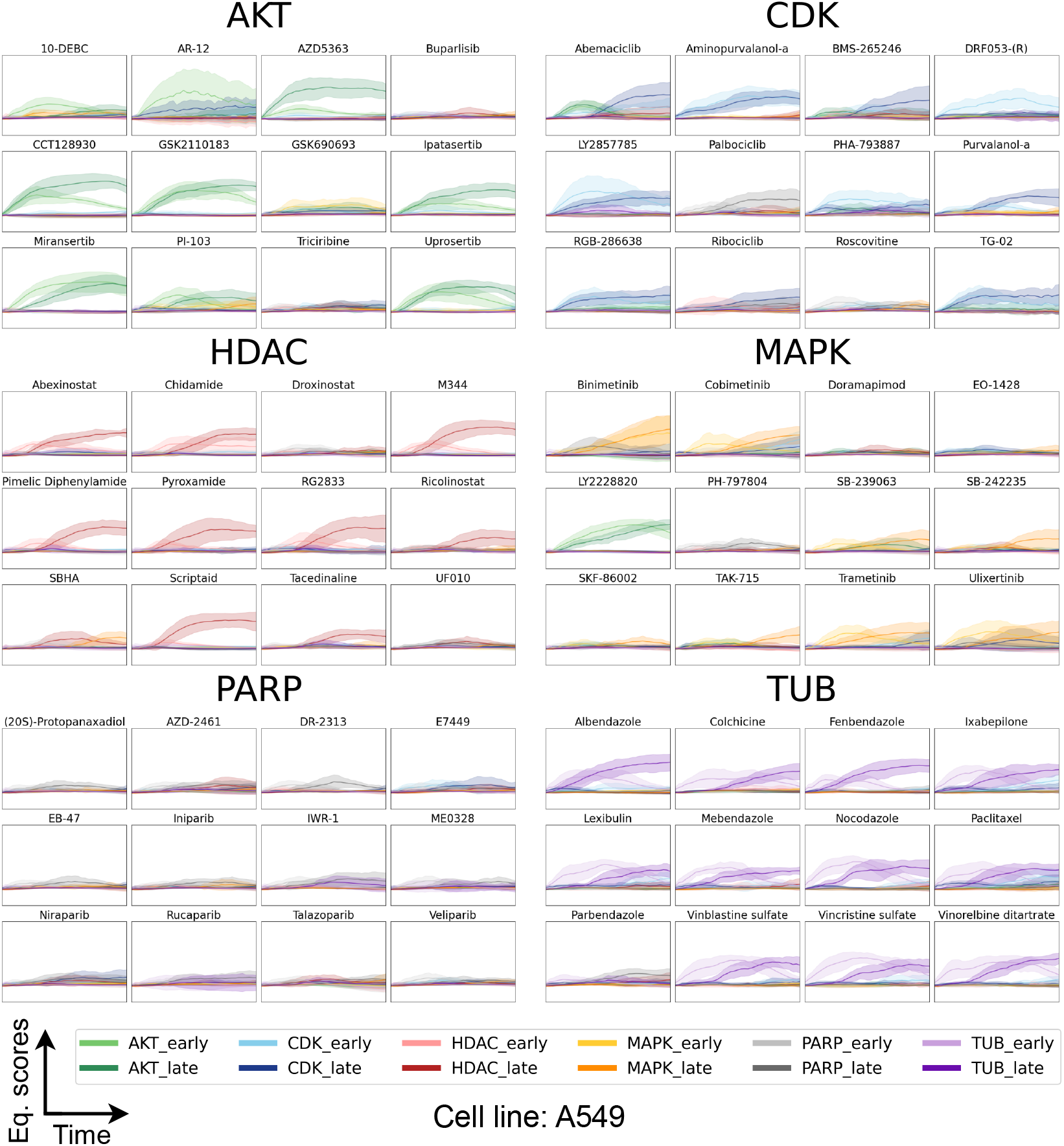
Eq. profiles for the compounds in the dataset grouped by their MOA. All axes have the same scale. The Y-axis represents the Eq. score predicted by the early and late models ranging from −0.5 to 2, while the X-axis shows time, ranging from 0 to 72 hours. Each line in the plot represents the mean Eq. profile from 15 wells across three plates, with one standard deviation depicted as the confidence interval. Consequently, each plot contains twelve Eq. profiles, with two profiles (early and late) for each MOA. Notably, in most cases, only 1-3 profiles are visible, as the others are predicted to be close to zero. Lighter colours indicate early Eq. profiles, whereas darker colours represent late Eq. profiles.

**Figure 6.**
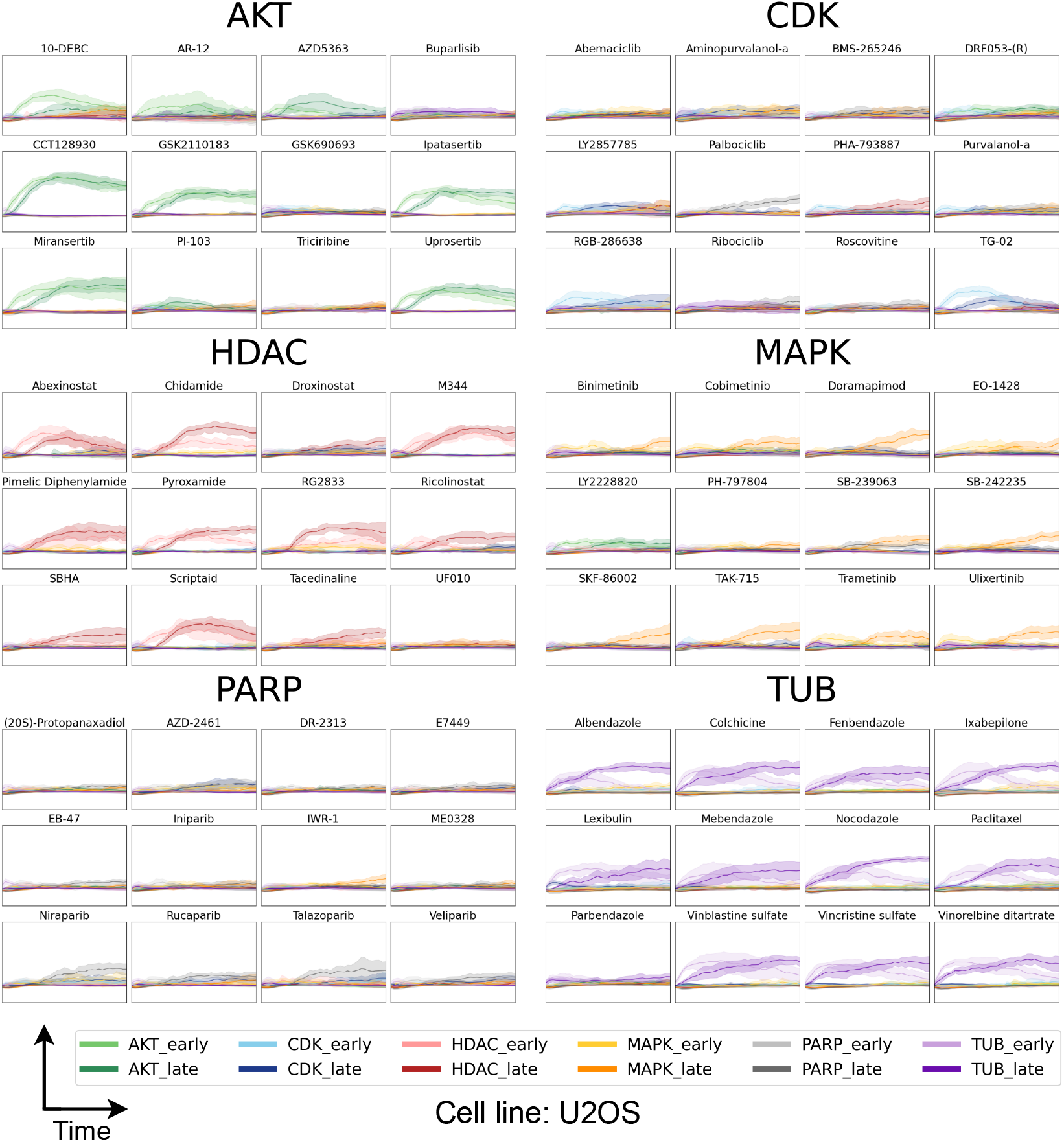
Eq. profiles for the compounds in the dataset grouped by their MOA. All axes have the same scale. The Y-axis represents the Eq. score predicted by the early and late models ranging from −0.5 to 2, while the X-axis shows time, ranging from 0 to 72 hours. Each line in the plot represents the mean Eq. profile from 15 wells across three plates, with one standard deviation depicted as the confidence interval. Consequently, each plot contains twelve Eq. profiles, with two profiles (early and late) for each MOA. Notably, in most cases, only 1-3 profiles are visible, as the others are predicted to be close to zero. Lighter colours indicate early Eq. profiles, whereas darker colours represent late Eq. profiles.

**Figure 7.**
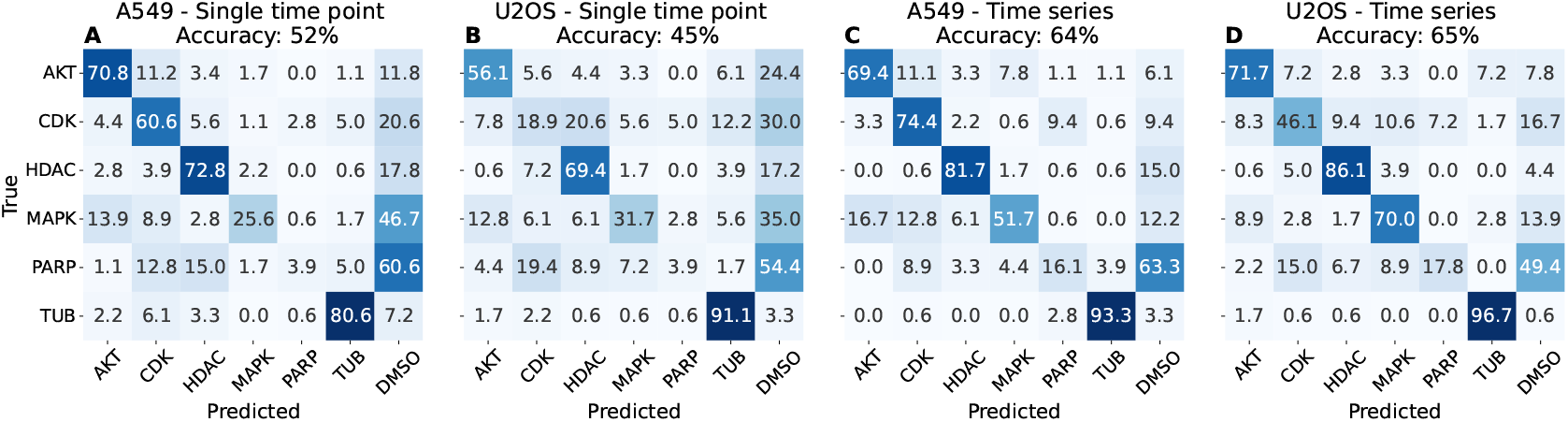
Confusion matrices of classification results using DINO features with the true labels on the Y-axis and the predicted on the X-axis. The data is normalized row-wise. Thus, each number represents the percentage of samples from a given true MOA that are classified into each predicted MOA group. In A and B, the results from a single time point (48h) classifications for A549- and U2OS cells with 52% and 45% accuracy respectively are shown. Both struggle with classifying MAPK and PAPR correctly and for U2OS, CDK is also hard. In C and D the results from time series classification for A549- and U2OS cells with 64% and 65% accuracy are shown. Here most MOAs are predicted clearly better compared to A and B. Especially MAPK going 25.6% accuracy in A to 51.7% in C for A549 cells and 31.7% in B to 70.0% in D. Observations are classified as negative controls (DMSO) if they are below the detection limit. Negative control observations are not included in the confusion matrix since they would by default be correct ~95% of the time due to the definition of the detection limit.

An example of a group that has clear, distinct early and late effects is the tubulin-targeting group (TUB compounds), which targets the cytoskeleton [35]. During mitosis, the cytoskeleton dynamically rearranges and this process is disrupted by treatment with these compounds resulting in an increase in mitotic index. This is observed at 8h in the image in Fig 4. This stage is followed by the cells presenting ruffled plasma membrane boundaries and an altered morphology distinct from untreated cells (see Fig 4). These two distinct morphologies are clearly seen in the Eq. profiles of Nocadazole (TUB) in Fig 4 for both cell lines with clear early and late profiles. In addition, small, rounded, mitotic cells are frequent in live-cell images taken at 8h but at a later time point, 48h, the cells exhibit a ruffled cell plasma membrane instead.

To investigate the temporal effects of different compounds, we generated CV Eq. profiles based on Eq. scores, smoothing, and a CV scheme. These profiles, grouped by MOA, are shown in Fig 5 for A549 and Fig 6 for U2OS.

From the Eq. profiles we can see that the AKT compounds exhibit no clear early-late distinction, instead showing high Eq. scores for both consistently throughout the majority of the time course. The major exceptions for AKT are Buparsilib, GSK690693, and Triciribine which all display low AKT Eq. scores in both cell lines (Fig 5 and 6). CDK compounds are overall more heterogeneous, yet like AKT, they lack a clear early-late distinction. What stands out are Abemaciclib and BMS-265246, demonstrating a notable early onset of AKT-late Eq. scores from hours 0 to 30. Additionally, DRF053-(R) is the only compound only showing an onset of CDK-early with no late profile.

Since the AKT and CDK pathways are both directly and indirectly connected, it has been shown that compounds inhibiting AKT (upstream) may indirectly reduce the activation of CDK 2, 4, and 6 (downstream) [36]. Additionally, the AKT-inhibitor AR-12 shows slight CDK-late Eq. scores in A549 cells. CDK4/6 inhibition has been previously reported to suppress AKT-mTOR signalling which might explain these findings [37].

Palbocicilib exhibit a PARP-late Eq. profile and Roscovitine exhibits little to no effect. In U2OS cells, most CDK compounds have low scores, with only LY2857785, RGB-286638, and TG-02 showing visible profiles. For HDAC compounds, clear late Eq. profiles along with less distinct early profiles are observed in A549 cells, Fig 5, with Droxinostat and UF010 showing lower scores than the rest. For U2OS cells both early and late HDAC-profiles are more clearly visible in Fig 6, with Droxinostat and UF010 showing low scores than the rest. For the MAPK compounds, the effects differ slightly between the two cell lines. While several compounds show both clear profiles in both, U2OS exhibits higher Eq. scores overall and more compounds. The exception being LY2228820 which display clear AKT-early and AKT-late profiles for both cell lines. PARP compounds generally present very subtle profiles, if any. For those showing a PARP profile, only late profiles are visible, consistent across both cell lines. TUB compounds display a clear early-late distinction in their Eq. profiles. With the exception of Parbendazole in both cell lines.

A supervised modelling-design like this reveals far more details about each compounds effect compared to an unsupervised such as UMAP (Fig 2F and G). However, when training supervised models, it must be done with care to avoid over-optimistic results and faulty conclusions. Therefore, to validate our models and findings, we consistently predict new compounds on new plates to mimic a real world drug screening scenario.

### 4.2. MOA classification - Time series vs single time point

Using the Eq. sums calculated from the Eq. profiles above, as well as the Eq. scores from models trained only on a single time point (48h), and their respective detection limits, we predict MOA for the compound-treated wells for models trained with the high-dimensional DINO features. For the MOA classification, each well is classified into one of seven classes (AKT, CDK, HDAC, MAPK, PARP, TUB, and DMSO). However, only wells belonging to the six MOA classes are classified and included in the accuracy calculations. Wells from the seventh class, DMSO i.e. negative controls, are not included, as this class would by default have an accuracy of ~95% based on the definition of the detection limit. Importantly, if a well’s highest Eq. sum falls below the detection limit, it is classified as DMSO. Additionally, we have performed the same analysis using various CV-schemes with DINO features. The different CV-schemes results in different accuracies but the trend between time series and single time point persists throughout (Supplementary information Fig S1). We have also included results from using different feature sets. We have compared features from VISTA2D [38, 39], cell properties calculated from the segmentation masks, and image statistics. The trend between time series and single time point also holds for all features sets, i.e. LCTP results in higher accuracy. These results are presented in the supplementary information.

Fig 7 shows the MOA classification accuracies from the DINO features for the single time point data and time series data using LCTP for the A549 and U2OS cells. In Fig 7A, the predictions for A549 cells at a single time point (48h) yield a total accuracy of 52%. AKT, CDK, HDAC, and TUB are classified with relatively high accuracy. Meanwhile, MAPK and more so PARP, often fail to pass the detection limit and are thus classified as negative controls (DMSO) with 46.7% of MAPK observations and 60.6% of PARP observations. Meaning, these were not significantly different from the negative controls.

For U2OS single time point classification (Fig 7B), the accuracy is lower at 45%, with increased misclassification across multiple MOAs. Most notably for CDK where fewer instances passes the detection limit and 30.0% ends up in the negative control class (DMSO) and 20.6% are predicted as HDAC. MAPK and PARP are again hard to distinguish from negative controls leading to 35.0% and 54.4% classified as DMSO respectively.

Time series data in combination with LCTP classification significantly improves accuracy for both cell lines across most MOAS with *p* = 0.0203 for A549 and *p* = 0.0088 for U2OS. In Fig 7C, the accuracy for A549 increases to 64%, with a pronounced reduction in MAPK misclassification into DMSO, from 25.6% correctly classified to 51.7%. Similarly, U2OS classification improves to 65%, with a noticeable increase for AKT, CDK, HDAC, and MAPK. Time series classification enhances overall performance, highlighting the advantage of incorporating temporal data for more accurate MOA classification, significantly improving accuracy for all MOAs for both cell lines.

The LCTP is highly modular serving two main purposes. First, it allows for easy substitution of models in each step when improvements become available. Segmentation-, live/dead classifying-, and feature extraction models are continuously improved, and as better models enter the scene, our proposed workflow will still be applicable and with, most likely, improved results. Second, it acts as a means of “concentrating” relevant information. Meaning, only the parts of the data we want to focus on are passed onto the next step. The segmentation model identifies the cells within images and by removing the background, we make sure the feature extraction model focus solely on the cells. Then, by only including the living cells based on the live/dead classification model, we limit the impact of cell death and focus on identifying more subtle morphological changes. Thus, we get more control over what goes into our median well profiles. Using these profiles, we train supervised, time-dependent models to create interpretable Eq. profiles that are both useful for temporal insights and MOA prediction with a thorough double-blind CV scheme. Doing the same CV scheme with an image-based classifier, i.e. a model that takes raw cell images as input and outputs a MOA label, would be considerably more computationally intensive, highlighting another benefit of our modular approach.

## 5. Conclusion

This paper introduces a novel workflow for analysing complex high-dimensional time series data from label-free live-cell imaging, yielding interpretable and biologically relevant results called LCTP. The workflow includes multiple pre-trained AI models to segment cells, classify them as live or dead, and extract high-dimensional single-cell features. These features serve as input to custom-trained ANN models, whose output reveals temporal drug responses, forming the basis for MOA classification.

Comparing MOA classification accuracy across different settings; single time point (48h) versus full time series demonstrates that combining time series data with high-dimensional features is optimal. A single time point may overlook transient states, while a full time series captures critical events essential for MOA prediction. Time series analysis also benefits from noise reduction through filtering, enhancing accuracy.

This underscores the notion that from label-free live-cell imaging, we can strategically extract information-rich features to predict the MOA of a compound. Beyond screening, this method is expected to be broadly useful for studying dynamic cellular responses, providing insight into how cells react to biological and chemical perturbations over time. It has potential to become one important component in phenotyping for next-generation drug discovery.

## Data and Code availability

All code and link to the dataset is available at https://github.com/edvinforsgren/LCTP.

## Declaration of generative AI and AI-assisted technologies in the manuscript preparation process

During the preparation of this work the author(s) used Chat-GPT in order to improve readability and grammar. After using this tool/service, the author(s) reviewed and edited the content as needed and take(s) full responsibility for the content of the published article.

