## Supplementary Information for "The time dimension matters: Improving mode of action classification with live-cell imaging"

### Supplementary Information for “The time dimension matters: Improving mode of action classification with live-cell imaging” - Artificial Intelligence in the Life Sciences

Edvin Forsgren[a], Jonne Rietdijk[b], David Holmberg[b], Julia Juneblad[c], Bianca Migliori[d], Martin M. Johansson[b], Jordi Carreras-Puigvert[b], Johan Trygg[c,e], Gillian Lovell[f], Ola Spjuth[b], and Pär Jonsson[c,e]*

[a] Sartorius BioAnalytics, Umeå, Sweden
[b] Department of Pharmaceutical Biosciences and Science for Life Laboratory, Uppsala University, Uppsala, Sweden
[c] Sartorius Corporate Research, Umeå, Sweden
[d] Sartorius Corporate Research, New York, USA
[e] Computational Life Science Cluster (CLiC), Department of Chemistry, Umeå University, Umeå, Sweden
[f] Sartorius BioAnalytics, Royston, UK

#### Comparing CV approaches

In the paper, we use a dual blind cross-validation scheme, meaning all predictions are done on unseen compound treatments from unseen plates. The alternatives are presented below where we do Compound wise (predicting wells with unseen compound treatments), plate wise (predicting wells from unseen plates), and 7-fold (randomly excluding every 7$^{th}$ well from the training set for each of the 7 rounds and predicting those). The accuracies are presented in Figure S1 below.

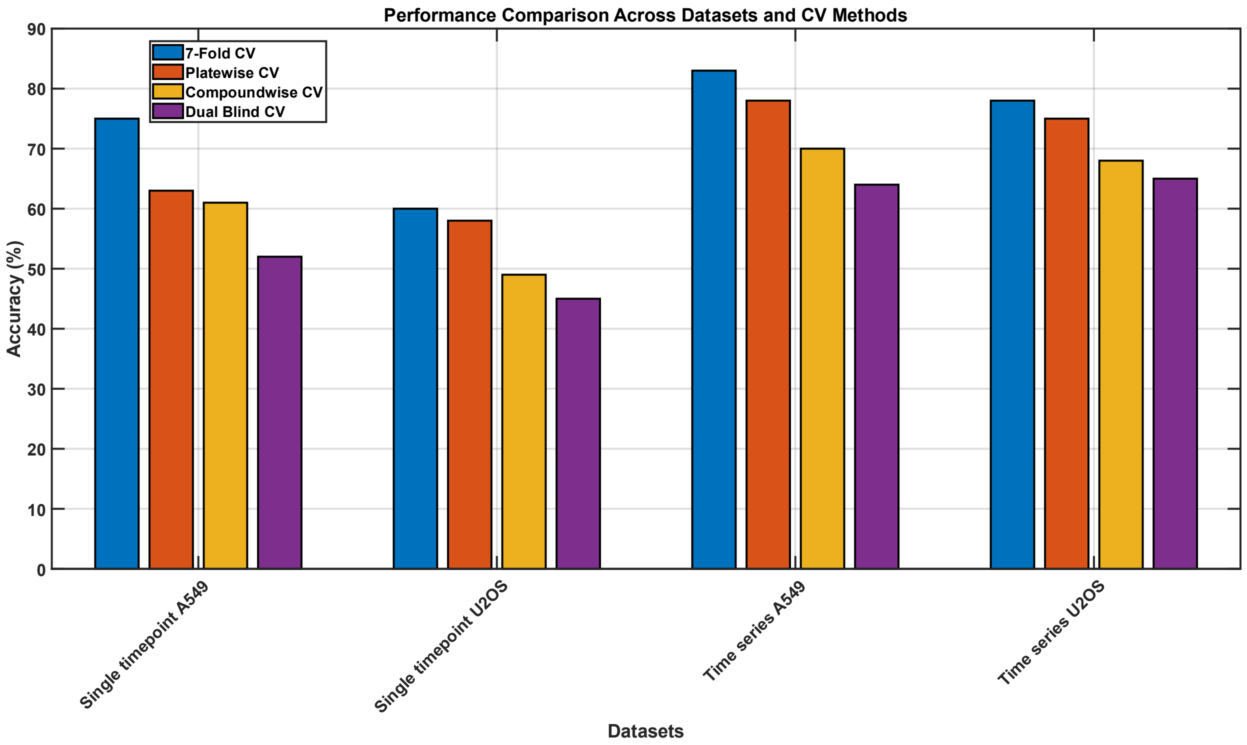


*Figure S1. Accuracies resulting from different CV-schemes. Blue is a 7-fold CV, orange is plate wise, yellow is compound wise, and purple is dual-blind CV.*

From Figure S1 we can see that the dual-blind CV is the most challenging setup since it the model is challenged to predict unseen compound treatments from unseen plates, simulating a real-world scenario. Noteworthy is that for all CV-schemes, our time series classification is more accurate than a classic single time point approach. The trend between the CV-schemes is also consistent for the different datasets, highlighting the importance of choosing the CV-scheme to not give over-optimistic results.

#### Distribution Plots of Cell Morphologies

To visualize differences in cell morphologies and biological variability between MOA groups and the two cell lines U2OS-WT and A549-WT, well-based distributions were compared for cell count, area, and eccentricity. The analysis presents both inter-cell line differences in Figure S2-S4, along with temporal changes within cell lines in Figure S5-S7. The y-axis is normalized within each MOA group for better comparison of distributions. The area represents number of pixels, the cell counts individual cells for each well, and eccentricity is the measure of how much an object deviates from a perfectly round circle, which would be 0. See Table S2 for calculation details.

#### Cell Type Comparison of Cell Properties

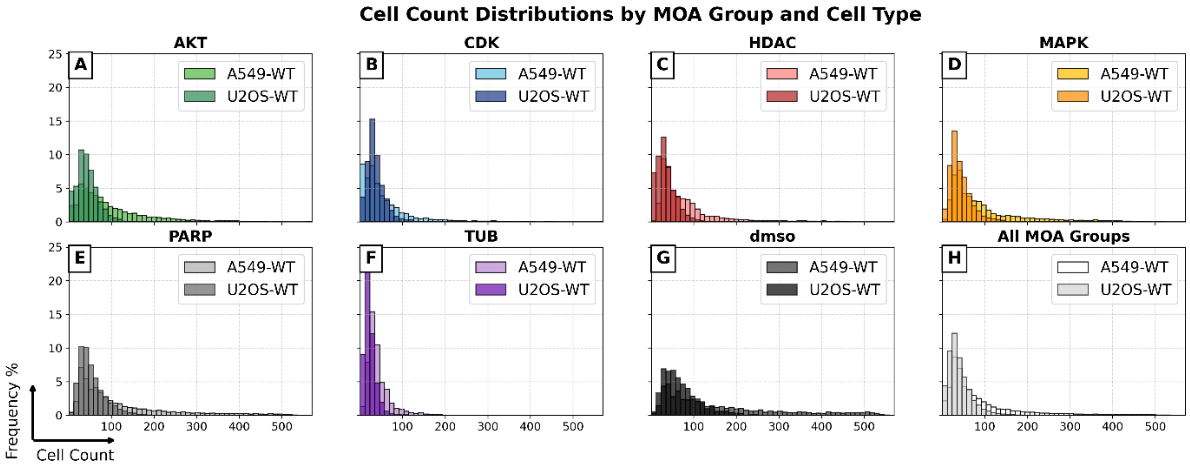


*Figure S2. Comparison between cell count in each well for the different MOA groups, untreaded negative controls (DMSO), and All MOA Groups. The A549-WT cell line has a slightly higher cell count than U2OS-WT. This is due to the proliferation rate being slightly higher for A549-WT. Additionally, CDK (B), HDAC (C), and TUB (F) have lower cell count than the other MOA groups.*

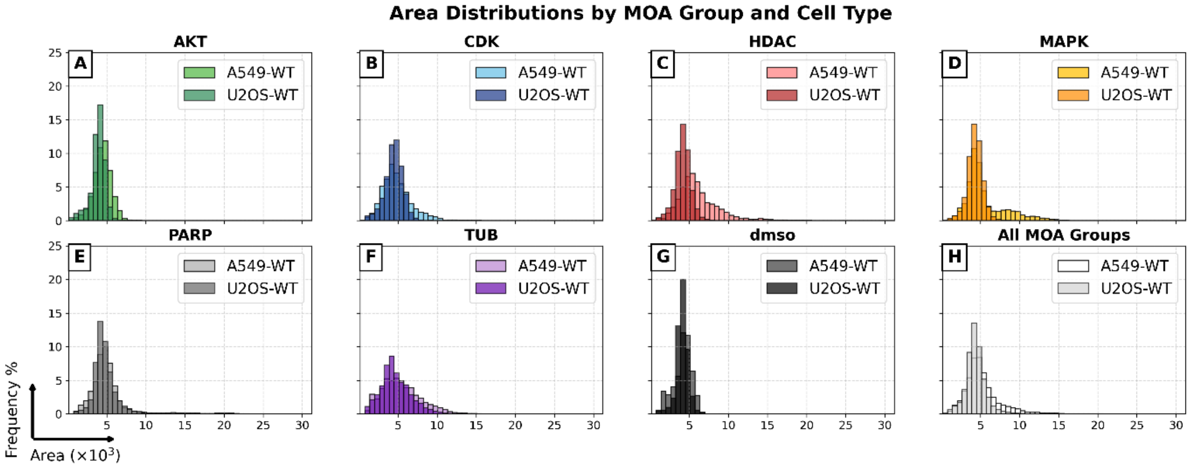


*Figure S3. Comparison between median cell area (pixels) in each well for the different MOA groups, untreaded negative controls (DMSO), and All MOA Groups. The A549-WT cell line has slightly larger cells than U2OS-WT. Additionally, HDAC (C) and MAPK (D) results in larger cells than the other MOA groups. This is also observed in the cell image examples presented in Figure 4 in the paper.*

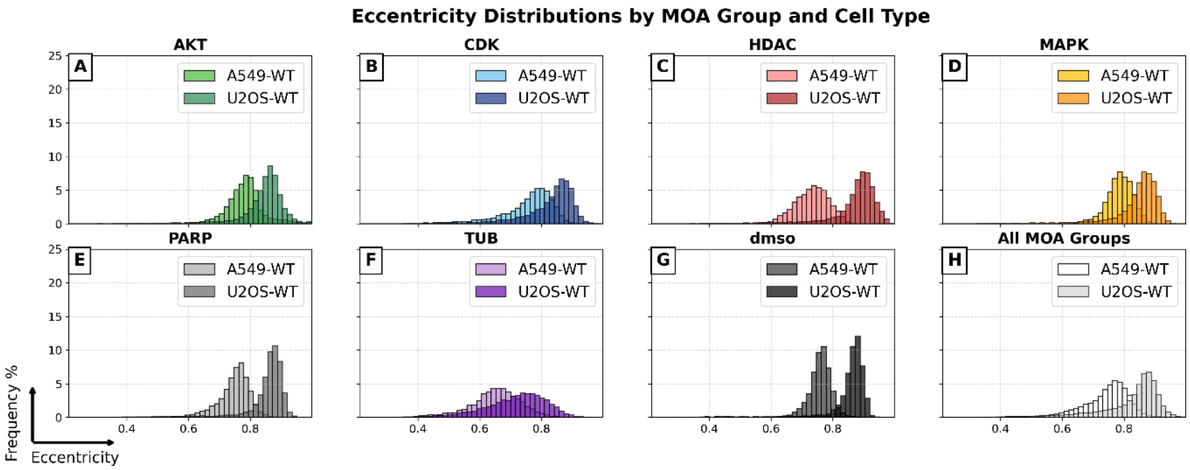


*Figure S4. Comparison between median cell eccentricity in each well for the different MOA groups, untreaded negative controls (DMSO), and All MOA Groups. The A549-WT cell line has lower eccentricity than U2OS-WT, meaning A549-WT cells are rounder. Additionally, AKT (A), CDK (B), MAPK (D), and TUB (F) result in more similar cells between the two cell lines than HDAC (C), PARP (E), and DMSO (G). Meaning the morphological changes induced by the compounds makes the cells more similar to each other in terms of eccentricity.*

#### Time Dependent Distributions of Cell Properties

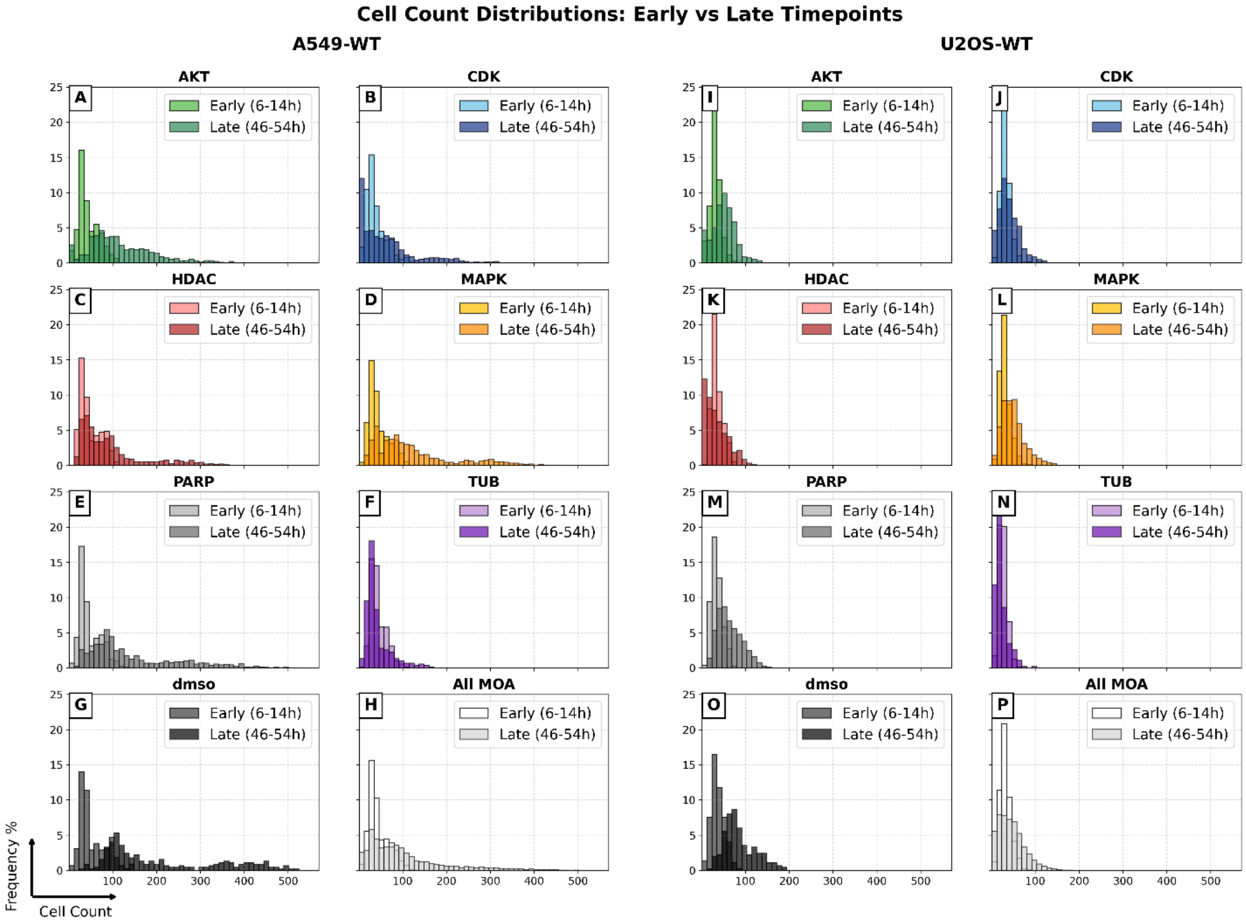


*Figure S5. Comparison of cell count between early and late time windows for A549-WT (A-H) and U2OS-WT (I-P). Noticeably, A549-WT has a higher cell count than U2OS-WT, especially for untreated negative controls (DMSO) (G and O) and lower effect compounds such as PARP (E and M) and MAPK (D and L). TUB (F and N) has a strong effect on cell proliferation since it hinders cells from exiting their mitotic state.*

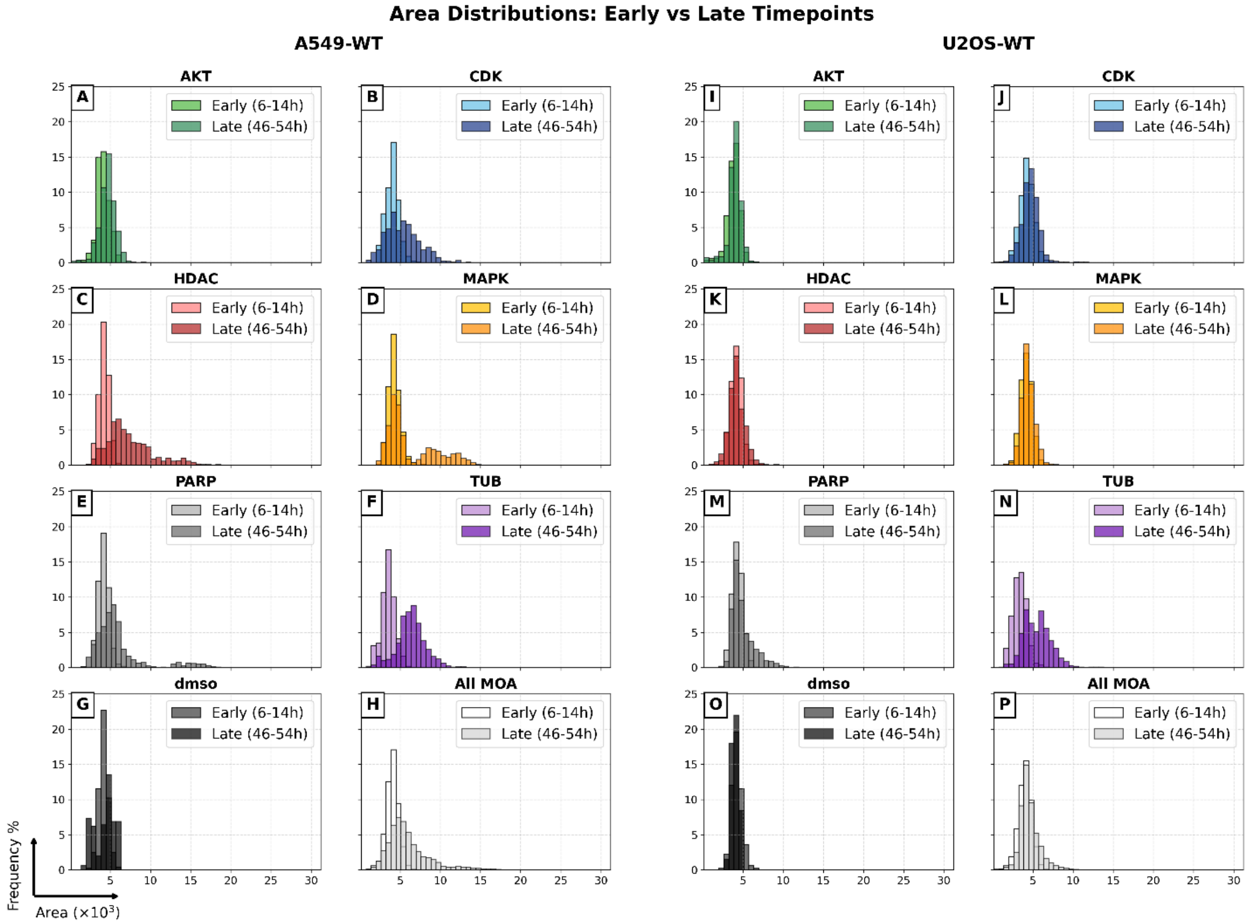


*Figure S6. Comparison of median Area between early and late time windows for A549-WT (A-H) and U2OS-WT (I-P). Noticeably, CDK (B and J), HDAC (C and K), and MAPK (D and L) induce a clear cell enlargement for late time points for A549-WT cells but not as clear for U2OS-WT. For A549-WT, MAPK (D) treatments are split into two distributions, one clear enlargement for late timepoints and one where the cell size remains largely the same as for early time points. This might explain why MAPK is harder to classify in the paper for A549-WT than U2OS-WT due to a more heterogeneous effect.*

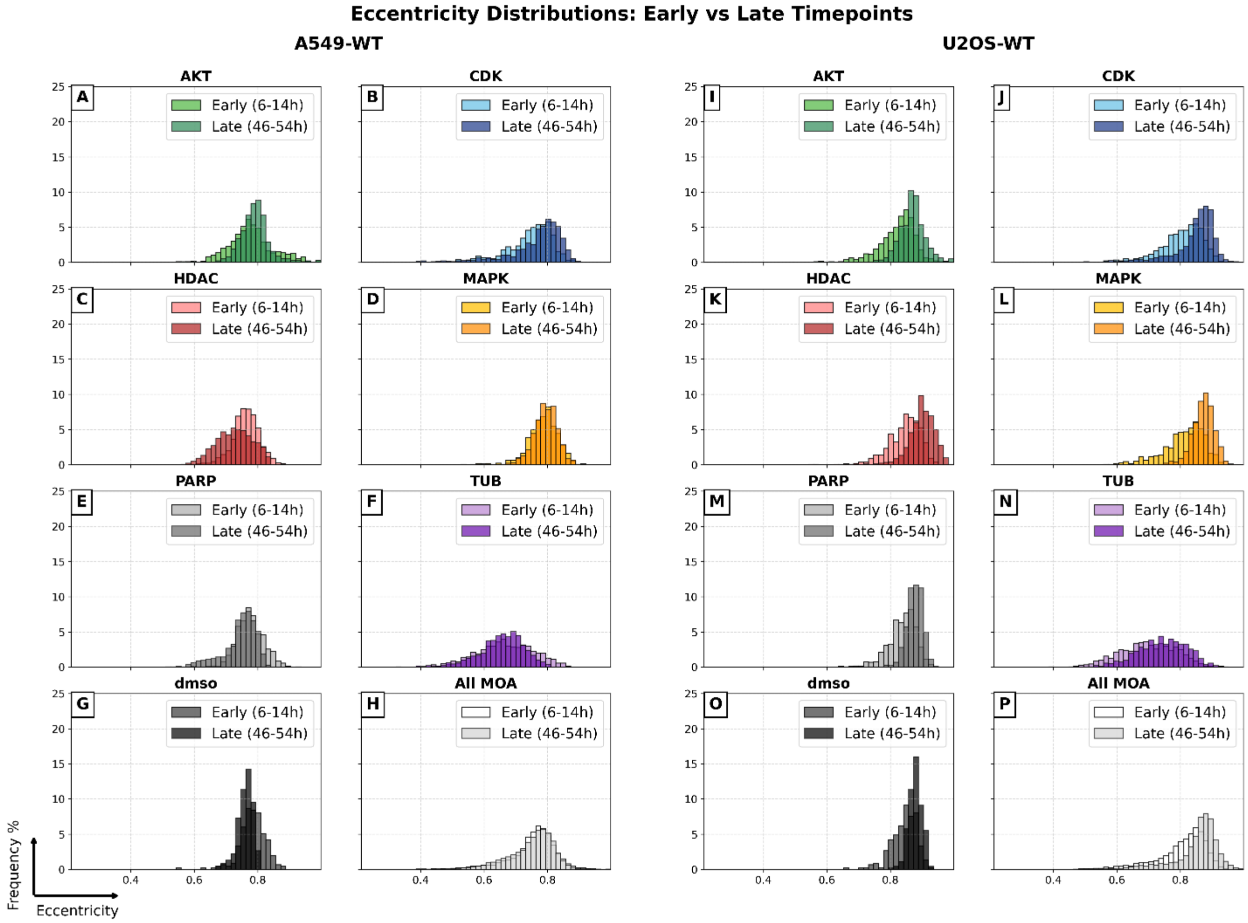


*Figure S7. Comparison of median Eccentricity between early and late time windows for A549-WT (A-H) and U2OS-WT (I-P). Noticeably, HDAC treatments on A549-WT (C) decrease the eccentricity of cells at late timepoints, i.e. making them rounder. The opposite happens for U2OS-WT cells treated with HDAC compounds (K), i.e. the eccentricity increases for later time points. Overall, U2OS-WT cells are more eccentric than A549-WT cells.*

#### LCTP Analysis using All Cells

In the paper we state:

*"We hypothesized that restricting the analysis to living cells would capture more relevant information about cellular condition and function, rather than reflecting post-mortem morphological changes."*

This is strengthened by the results presented in Figure S8.

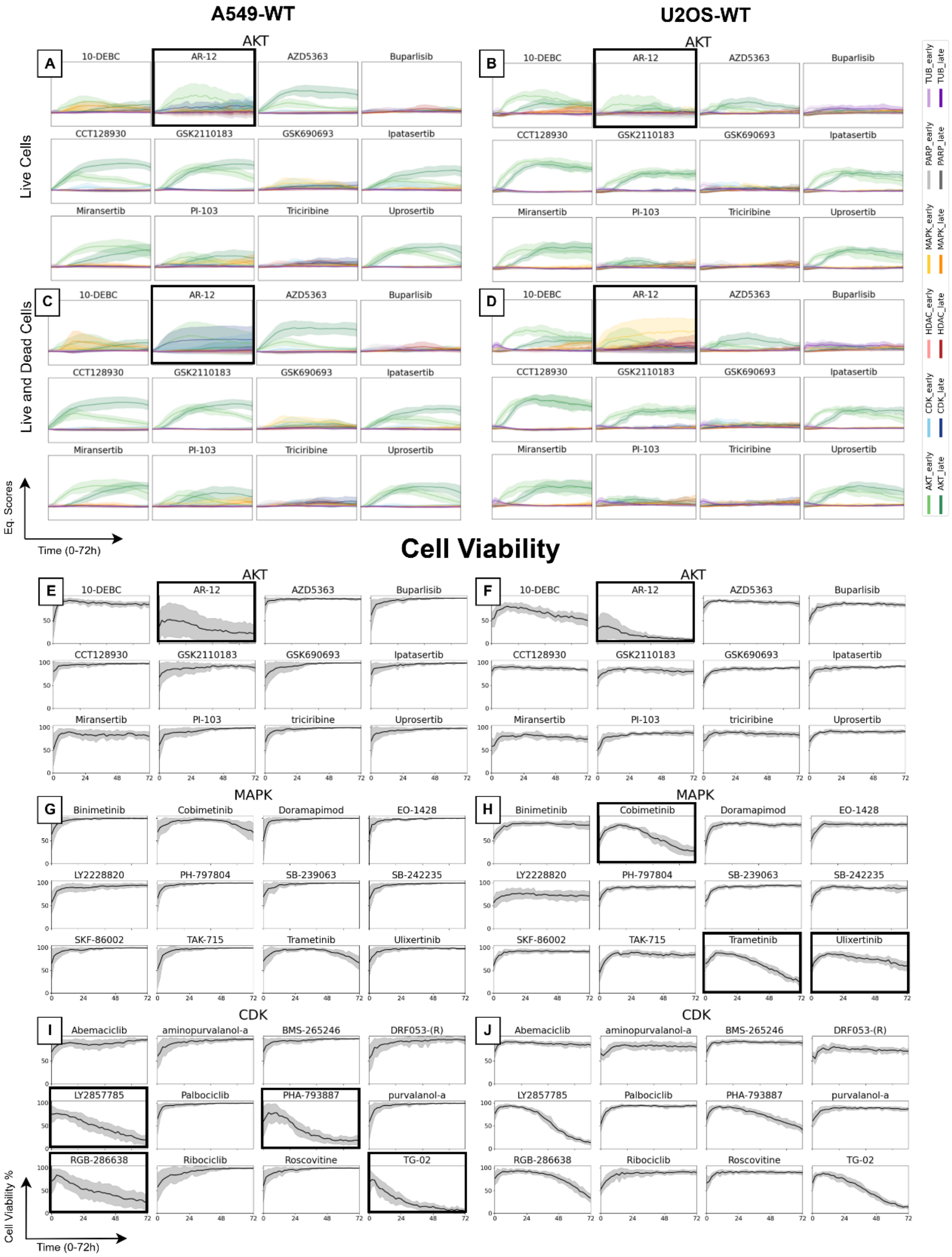


*Figure S8. The two first rows show the CV Eq. Profiles for AKT inhibiting compounds. The left column (A, C, E, G, and I) shows results from the dataset with A549-WT cells and the right column (B, D, F, H, and J) shows results from the dataset with U2OS-WT cells. In A and B, only living cells were included to form the median well features and in C and D both living and dead cells were included. E-J shows the mean viability over time for compounds from different modes of action (MOA) groups. The main takeaway from this is that when dead cells are included, AR-12 (AKT inhibiting compound resulting in lower viability, E and F) is confounded with either CDK or MAPK (C and D) for the different cell types. Looking at the viability plots for the different cell types for CDK and MAPK (G-J), we can see a difference in the viability between the cell types which correlate with the results in C and D.*

Here, A and B are the Eq. scores for AKT from a median-well feature set resulting from only living cells (the approach we chose in the paper). In C and D, we have included both living and dead cells in the median well calculation. Notably, the AR-12 compound is predicted as CDK for A549-WT cell line and as MAPK for U2OS-WT cell line. Looking more in depth into the cell viability for AKT, CDK, and MAPK (E-J) we first note that AR-12 has considerably lower viability (high cell death) than the rest of the AKT-compounds (E and F). Next, we can see that MAPK differs significantly between the two cell lines. For A549-WT (G), the viability is generally very high but for U2OS-WT (H) there are a few compounds (highlighted in H) that have considerably lower viability. Comparing the same difference between the cell lines for CDK (I and J) we can see that both have compound treatments inducing cell death but resulting in lower viability for A549-WT (I).

In summary, AR-12 (having lower viability than the rest of AKT-inhibiting compounds) is predicted as CDK for A549-WT and MAPK for U2OS-WT when dead cells are included but not when we only focus on living cells. Meaning that when we include dead cells, the dead cell morphology will affect the features and confound the models to draw focus from the actual, more subtle morphological changes that reflect the phenotype induced from the mechanism of action.

In Figure S9, we can see that U2OS time series (D) has slightly lower accuracy than when only including living cells and that AKT are more commonly misclassified as MAPK than in the paper (Figure 7). Otherwise, overall accuracy remains at a similar level as the results presented in the paper.

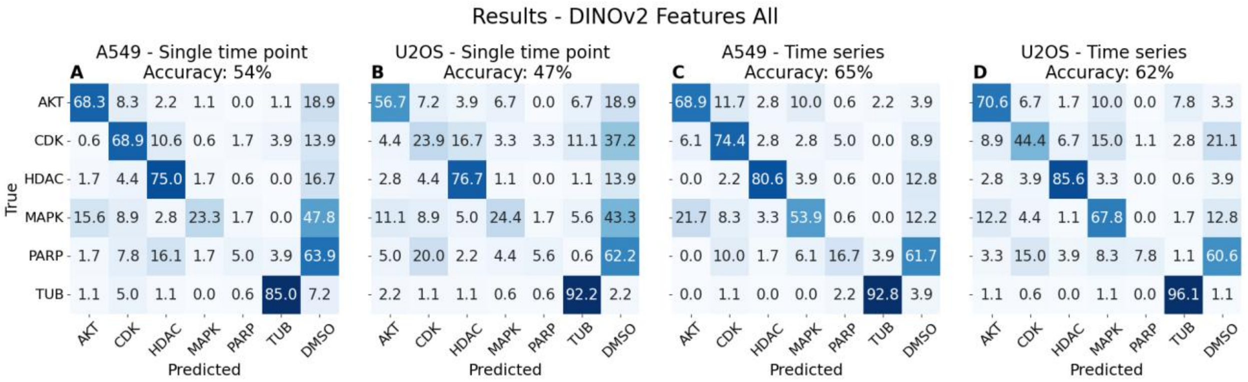


*Figure S9. Confusion matrices of classification results using DINO features from both living and dead cells in the median well aggregation, with the true labels on the Y-axis and the predicted on the X-axis. The data is normalized row-wise. Thus, each number represents the percentage of samples from a given true MOA that are classified into each predicted MOA group.*

#### MOA Classification with LCTP – Feature set Comparison

##### Image Statistics Features

We extracted 33 image statistics properties on the whole image, using segmentation masks to separate foreground and background statistics. These statistics are presented in Table S1 and were calculated with standard numerical operations and scikit-image filters module [3]. In addition to training all image statistics features, training was performed using background and foreground statistics separately, where foreground features represent the cell regions. Although both foreground and background statistics show low accuracy, the foreground presented a slightly higher accuracy out of the two. This concludes that the MOA classification is not only picking up image specific features from the background. The ANNs trained and used for prediction contain two layers consisting of 8 nodes each; other parameters stayed the same as in the paper. Results are presented in Figure S10.

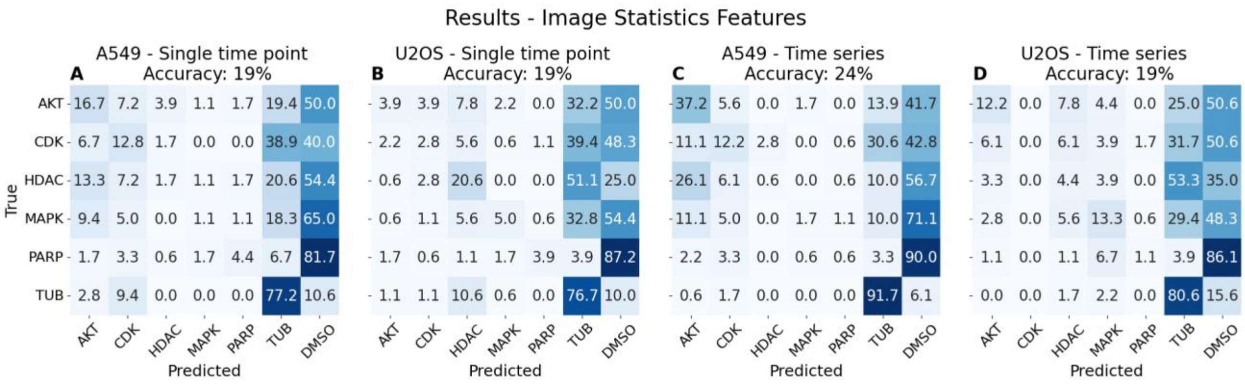


*Figure S10. Confusion matrices of classification results using image statistic features, with the true labels on the Y-axis and the predicted on the X-axis. The data is normalized row-wise. Thus, each number represents the percentage of samples from a given true MOA that are classified into each predicted MOA group.*

*Supplementary Table S1. Summary of image statistics features.*

| Whole image Intensity Features | Texture and Edge Features | Foreground and background Features |
|---|---|---|
| Minimum Intensity | Granularity | Mean Intensity |
| Maximum Intensity | Laplacian Variance | Standard Deviation of Intensity |
| Mean Intensity | Sobel Mean | 95th Percentile Intensity |
| Standard Deviation of Intensity | Sobel Standard Deviation | Granularity |
| 1st Percentile Intensity | Edge Density | Sobel Mean |
| 5th Percentile Intensity | RMS Contrast | Sobel Standard Deviation |
| 95th Percentile Intensity | Entropy | Entropy |

| | | |
|---|---|---|
| 99th Percentile Intensity | | Foreground Fraction |
| Interquartile Range (IQR) of Intensity | | |
| Coefficient of Variation | | |
| Dynamic Range | | |

#### VISTA2D Features

Using VISTA2D [1, 2] model weights provided by NVIDIA (CC-BY-NC-SA-4.0 license, weight by request), we extracted 256 VISTA2D features based on the segmentation masks from AI Cell Health® from which we aggregated median well features. The ANNs trained and used for prediction contain two layers consisting of 256 and 128 nodes, respectively; other parameters stayed the same as in the paper. Results are presented in Figure S11.

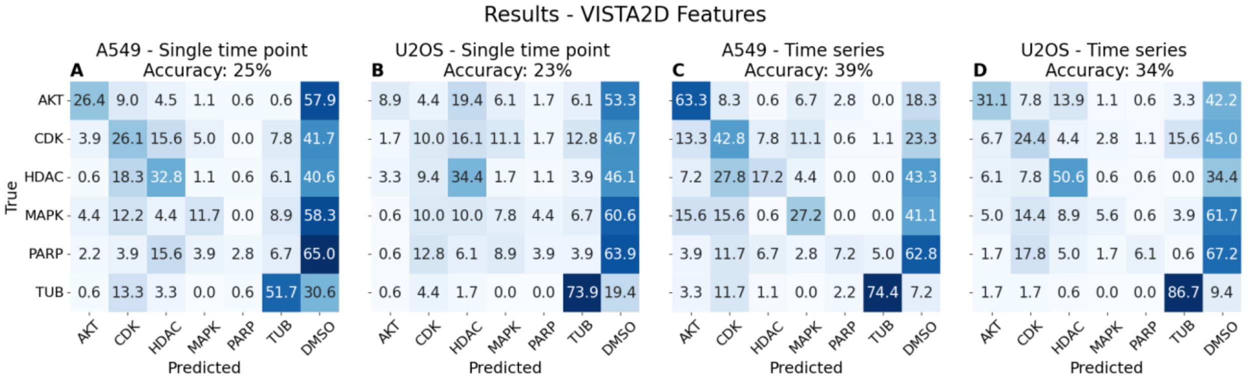


*Figure S11. Confusion matrices of classification results features extracted with VISTA2D, with the true labels on the Y-axis and the predicted on the X-axis. The data is normalized row-wise. Thus, each number represents the percentage of samples from a given true MOA that are classified into each predicted MOA group.*

#### Cell Properties extracted with scikit-image regionprops

We extracted 17 cell properties using scikit-image regionprops and SciPy stats [3, 4] based on the segmentation masks from AI Cell Health® from which we aggregated median well features. The features are presented in Table S2. The ANNs trained and used for prediction contain two layers consisting of 8 nodes each; other parameters stayed the same as in the paper. Results are presented in Figure S12.

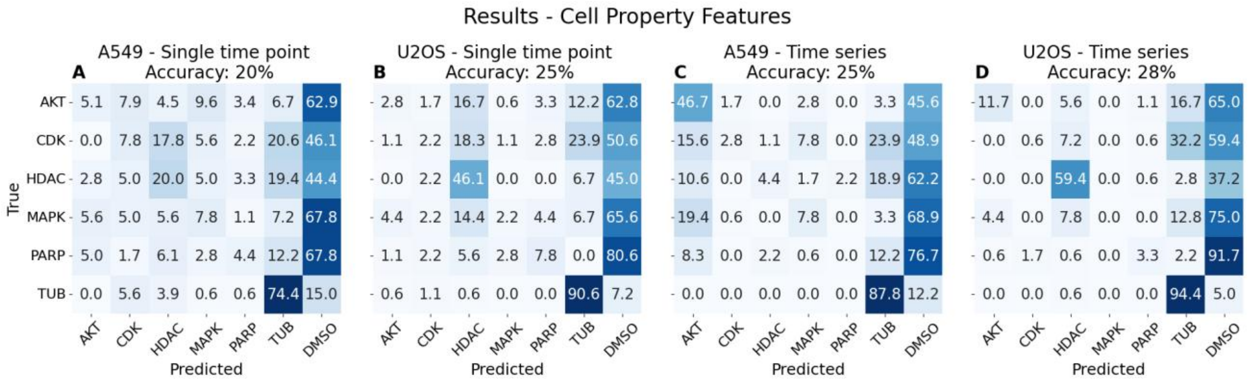


*Figure S12. Confusion matrices of classification results using features extracted with scikit-image regionprops based on the cell segmentation masks, i.e. basic single cell properties. The true labels on the Y-axis and the predicted on the X-axis. The data is normalized row-wise. Thus, each number represents the percentage of samples from a given true MOA that are classified into each predicted MOA group.*

*Supplementary Table S2. Summary of Cell Property features.*

| **Feature** | **Extraction Method** | **Unit/range** |
|---|---|---|
| Area | Scikit-image regionprops | pixels |
| Eccentricity | Scikit-image regionprops | 0-1 |
| Solidity | Scikit-image regionprops | 0-1 |
| Perimeter | Scikit-image regionprops | pixels |
| Feret | Scikit-image regionprops | pixels |
| Aspect Ratio | Scikit-image regionprops | ≥1 |
| Roundness | Area / (π × major_axis_length$^2$) | 0-1 |
| Circularity | 4π × Area / Perimeter$^2$ | 0-1 |
| Intensity (mean, max, min, std) | Scikit-image regionprops | arbitrary |
| Skewness, kurtosis | SciPy stats | dimensionless |
| Percentile (5, 95) | SciPy stats | arbitrary |
| Centroid Diff | √((centroid – weighted_centroid)$^2$) | pixels |

#### Model Accuracy Comparison

Table S3 summarizes the compared method accuracies, none of which are close to the DINOv2 accuracies in the paper. Meaning DINOv2 generates detailed and biologically relevant features suitable for MOA classification.

*Supplementary Table S3. Summary of accuracy with different feature sets.*

| Feature set | Cell type | Single Time Point | Time Series |
|---|---|---|---|
| **Image Statistics** | A549-WT | 19% | 24% |
| | U2OS-WT | 19% | 19% |
| **Vista2d (NVIDIA Model Weight)** | A549-WT | 25% | 39% |
| | U2OS-WT | 23% | 34% |
| **Cell properties** | A549-WT | 20% | 25% |
| | U2OS-WT | 25% | 28% |

### Consumables table

*Supplementary Table S4. Reagent and Consumable Details*

| Product | Supplier | Cat. Number | Concentration |
|---|---|---|---|
| DMEM (U2-OS-WT cells) | Gibco | 11965092 | |
| F12K (A549-WT cells) | Gibco | 21127022 | |
| FBS | Hyclone | SH30071.03 | 10% |
| Pen/strep | Gibco | 15140-122 | 1% |
| PBS | Gibco | 14190 | |
| Trypsin-EDTA | Gibco | 25200 | 0.25% |

### AI Cell Health image analysis parameters

*Supplementary Table S5. AI Cell Health Image analysis Parameters*

| AI Cell Health Analysis | A549 | U2OS |
|---|---|---|
| Segmentation sensitivity | 0.5 | 0.6 |
| Live/ Dead Classification threshold | 0.2 | 0.2 |
| Area filter | None | Minimum 100 $\mu m^2$ Maximum 2500 $\mu m^2$ |

[4] Pauli Virtanen, Ralf Gommers, Travis E. Oliphant, Matt Haberland, Tyler Reddy, David Cournapeau, Evgeni Burovski, Pearu Peterson, Warren Weckesser, Jonathan Bright, Stéfan J. van der Walt, Matthew Brett, Joshua Wilson, K. Jarrod Millman, Nikolay Mayorov, Andrew R. J. Nelson, Eric Jones, Robert Kern, Eric Larson, CJ Carey, İlhan Polat, Yu Feng, Eric W. Moore, Jake VanderPlas, Denis Laxalde, Josef Perktold, Robert Cimrman, Ian Henriksen, E.A. Quintero, Charles R Harris, Anne M. Archibald, Antônio H. Ribeiro, Fabian Pedregosa, Paul van Mulbregt, and SciPy 1.0 Contributors. SciPy 1.0: Fundamental Algorithms for Scientific Computing in Python. Nature Methods, 17(3), 261-272, 2020. DOI: 10.1038/s41592-019-0686-2.
